# Family-based quantitative trait meta-analysis implicates rare noncoding variants in *DENND1A* in pathogenesis of polycystic ovary syndrome

**DOI:** 10.1101/460972

**Authors:** Matthew Dapas, Ryan Sisk, Richard S. Legro, Margrit Urbanek, Andrea Dunaif, M. Geoffrey Hayes

**Affiliations:** Division of Endocrinology, Metabolism, and Molecular Medicine, Department of Medicine, Northwestern University Feinberg School of Medicine, Chicago, IL; Department of Obstetrics and Gynecology, Penn State College of Medicine, Hershey, PA; Center for Genetic Medicine, Northwestern University Feinberg School of Medicine, Chicago, IL; Center for Reproductive Science, Northwestern University Feinberg School of Medicine, Chicago, IL; Division of Endocrinology, Diabetes and Bone Disease, Icahn School of Medicine at Mount Sinai, New York, NY; Department of Anthropology, Northwestern University, Evanston, IL

## Abstract

Polycystic ovary syndrome (PCOS) is among the most common endocrine disorders of premenopausal women, affecting 5-15% of this population depending on the diagnostic criteria applied. It is characterized by hyperandrogenism, ovulatory dysfunction and polycystic ovarian morphology. PCOS is a leading risk factor for type 2 diabetes in young women. PCOS is highly heritable, but only a small proportion of this heritability can be accounted for by the common genetic susceptibility variants identified to date. To test the hypothesis that rare genetic variants contribute to PCOS pathogenesis, we performed whole-genome sequencing on DNA from 62 families with one or more daughters with PCOS. We tested for associations of rare variants with PCOS and its concomitant hormonal traits using a quantitative trait meta-analysis. We found rare variants in *DENND1A* (*P*=5.31×10^−5^, *P_adj_*=0.019) that were significantly associated with reproductive and metabolic traits in PCOS families. Common variants in *DENND1A* have previously been associated with PCOS diagnosis in genome-wide association studies. Subsequent studies indicated that *DENND1A* is an important regulator of human ovarian androgen biosynthesis. Our findings provide additional evidence that *DENND1A* plays a central role in PCOS and suggest that rare noncoding variants contribute to disease pathogenesis.

## INTRODUCTION

Polycystic ovary syndrome (PCOS) is a common endocrine disorder affecting 5-15% of premenopausal women worldwide^1^, depending on the diagnostic criteria applied. PCOS is diagnosed by two or more of its reproductive features of hyperandrogenism, ovulatory dysfunction, and polycystic ovarian morphology. It is frequently associated with insulin resistance and pancreatic β-cell dysfunction, making it a leading risk factor for type 2 diabetes in young women^2^.

PCOS is a highly heritable complex genetic disorder. Analogous to other complex traits^3^, common susceptibility loci identified in genome-wide association studies (GWAS)^4–8^ account for only a small proportion of the estimated genetic heritability of PCOS^9^. As GWAS were designed to assess common allelic variants, usually with minor allele frequencies (MAF) ≥2-5%, it has been proposed that less frequently occurring variants with greater effect sizes account for the observed deficit in heritability^10^. Next generation sequencing approaches have identified rare variants that contribute to complex disease pathogenesis^11–16^.

We tested the hypothesis that rare variants contribute to PCOS by conducting family-based association analyses using whole-genome sequencing data. We filtered and weighted rare variants (MAF ≤2%) according to their predicted levels of deleteriousness and grouped them regionally and by genes. We not only tested for associations with PCOS diagnosis, but also with its correlated, quantitative reproductive and metabolic trait levels: testosterone (T), dehydroepiandrosterone sulfate (DHEAS), luteinizing hormone (LH), follicle stimulating hormone (FSH), sex hormone binding globulin (SHBG), and fasting insulin (I). We then combined the quantitative trait phenotypes using a meta-analysis approach to identify sets of rare variants that associate with altered hormonal levels in PCOS.

## SUBJECTS AND METHODS

### Subjects

This study included 261 individuals from 62 families with PCOS who were Caucasians of European ancestry. Families were ascertained by an index case who fulfilled the National Institutes of Health (NIH) criteria for PCOS^17^. Each family consisted of at least a proband-parent trio. Brothers were not included in this study. Phenotypic data and some genetic analyses on these subjects have been previously reported^18–20^. The study was approved by the Institutional Review Boards of Northwestern University Feinberg School of Medicine, Penn State Health Milton S. Hershey Medical Center, and Brigham and Women’s Hospital. Written informed consent was obtained from all subjects prior to study.

Phenotyping for the dichotomous trait analysis (affected vs. unaffected) was performed as previously described^21^. Women were ages 14-63 years, in good health and not taking medications known to alter reproductive or metabolic hormone levels for at least one month prior to study. They had each had both ovaries and a uterus. Exogenous gonadal steroid administration was discontinued at least three months prior to the study. Thyroid, pituitary, and adrenal disorders were excluded by appropriate tests^21^. Women were considered to be of reproductive age if they were between the ages of at least 2 years post-menarche and 45 years old, and had FSH levels ≤40mIU/mL. Hyperandrogenemia was defined by elevated levels of T (>58 ng/dL), non-SHBG bound T (uT; >15 ng/dL), and/or DHEAS (>2683 ng/mL). Ovarian dysfunction was defined as ≤6 menses per year. Ovarian morphology was not assessed because it does not correlate with the endocrine phenotype^6, 22^.

Reproductive-age women with hyperandrogenemia and ovarian dysfunction were assigned a PCOS phenotype. Reproductive-age women with normal androgen levels and regular menses (every 27-35 days) were assigned an unaffected phenotype. Reproductive-age women with hyperandrogenemia and regular menses were assigned a hyperandrogenemic (HA) phenotype. Because androgen levels do not decrease during the menopausal transition^23^, women between 46-63 years with HA were also assigned the HA phenotype, regardless of menstrual cycle pattern. One index case fulfilled the criteria for PCOS when she was 45 years but she was 46 years when enrolled in the study. She was confirmed to have persistent HA and ovarian dysfunction. As done in our previous linkage^22^ and family-based association testing^24^ studies, women with both PCOS and HA phenotypes were considered affected. In the present study, we also included HA women between 46-63 years as affected. Women with normal androgen levels who were not of reproductive age and all fathers in the study were not assigned a phenotype.

The quantitative trait analysis examined associations between rare variants and T, DHEAS, SHBG, LH, FSH and insulin levels. In addition to the women included in the dichotomous trait analysis, women were included for quantitative trait association testing as follows. No additional women were included in the LH or FSH analyses. Women 46-72 years old were included in the analyses for T and DHEAS since androgen levels do not change during the menopausal transition^23^. These women were also included in the analysis of SHBG and insulin. Women with bilateral oophorectomy (n=10) not receiving postmenopausal hormone therapy were included in the analysis of the adrenal androgen, DHEAS, and insulin^25, 26^. We compared hormonal traits in women receiving postmenopausal hormone therapy (n=15) to women from the cohort of comparable age who were not on receiving hormonal therapy (n=10). Only SHBG levels differed significantly. Therefore, women receiving postmenopausal hormone therapy were included in the analyses of T and DHEAS. They were not included in the insulin analysis because of the effect of estrogen on circulating insulin levels^27^.

Fathers were included in the insulin level association test since there are not sex differences in this parameter^28^. Subjects receiving glucocorticoids (men=0, women=2) were excluded from all quantitative trait association tests. Where applicable, subjects receiving anti-diabetic medications (men=6, women=7) were excluded from the T, SHBG, and insulin analyses but were included in the DHEAS analysis^29^. Subjects with type 2 diabetes not receiving medications (men=3, women=7) were excluded from the insulin analysis but the women were included in the T, DHEAS and SHBG analyses.

Reference ranges for hormonal parameters from concurrently studied reproductively normal control subjects of comparable age, sex, BMI and ancestry^6, 30, 31^ are included in **Table 1**. All control subjects had normal glucose tolerance with a 75g oral glucose tolerance test^32^. Reproductive age control women were 18-45 years with regular menses, FSH<40 mIU/ml and normal androgen levels. Older control women were 46-65 years with a history of regular menses and normal androgen levels. Control men were 46-65 years.

**Table 1.** Clinical and biochemical characteristics of study population.

| Subject Type | Age<br>(years)<sup>a</sup> | BMI<br>(kg/m<sup>2</sup>) | Testosterone<br>(ng/dL) | SHBG<br>(nmol/L) | DHEAS<br>(ng/mL) | Insulin<br>(μU/mL) | LH<br>(mIU/mL) | FSH<br>(mIU/mL) |
| --- | --- | --- | --- | --- | --- | --- | --- | --- |
| **Index Cases** | 62 29 (19-48) | 62 35.6 (27.9-42.7) | 62 74 (65-90) | 59 56 (36-92) | 61 2095 (1638-2710) | 62 21 (14-29) | 59 14 (7-20) | 59 9 (8-11) |
| **Fathers** | 62 58 (43-85) | 61 28.9 (27.0-32.2) | - | - | - | 49 17 (11-26) | - | - |
| **Mothers** |  |  |  |  |  |  |  |  |
| HA | 6 51 (40-63) | 6 34.5 (25.2-38.2) | 5 51 (49-72) | 5 117 (105-135) | 6 2630 (1264-2694) | 3 19 (15-22) | - | - |
| Unaffected | 2 44 (42-45) | 2 24.0 (20.6-27.4) | 2 32 (27-37) | 2 280 (218-342) | 2 1081 (901-1260) | 2 12 (8-15) | 2 7 (6-7) | 2 8 (6-10) |
| Over 45 yo | 54 57 (46-72) | 53 28.6 (26.6-35.5) | 36 26 (16-34) | 25 93 (74-166) | 50 645 (501-993) | 28 19 (11-25) | - | - |
| **Sisters** |  |  |  |  |  |  |  |  |
| PCOS | 10 29 (19-36) | 10 32.3 (29.0-37.3) | 10 63 (49-74) | 10 55 (45-104) | 10 1665 (807-2774) | 10 28 (23-32) | 10 10 (5-15) | 10 11 (10-12) |
| HA | 5 32 (15-40) | 5 22.3 (22.3-28.5) | 5 67 (54-68) | 5 126 (69-177) | 5 2047 (1509-3775) | 5 15 (11-17) | 4 4 (3-18) | 4 10 (7-14) |
| Unaffected | 57 32 (14-45) | 57 25.1 (22.2-29.3) | 57 30 (23-35) | 55 128 (90-181) | 57 1408 (1028-1814) | 55 12 (9-15) | 55 5 (3-10) | 55 10 (7-12) |
| Over 45 yo | 3 47 (46-49) | 3 26.4 (24.5-32.1) | 3 30 (20-48) | 3 137 (88-150) | 3 772 (765-1588) | 3 13 (12-15) | - | - |
| **Reference Ranges** |  |  |  |  |  |  |  |  |
| Reproductive Aged Women | 346 30 (25-35) | 346 28.5 (23.0-35.4) | 227 29 (21-37) | 188 100 (70-144) | 226 1357 (1018-1756) | 185 12 (10-16) | 173 4 (3-8) | 173 9 (7-12) |
| Men | 55 53 (50-57) | 55 28.0 (25.9-31.0) | - | - | - | 46 14 (10-17) | - | - |
| Older Women | 69 56 (52-60) | 69 26.4 (23.2-29.6) | 69 23 (15-30) | 60 102 (75-156) | 66 645 (487-1016) | 69 13 (10-17) | 48 50 (35-69) | 49 80 (57-100) |

Traits reported as count, median (25<sup>th</sup>-75<sup>th</sup> percentiles). Values reported here are unadjusted. Trait values were adjusted and normalized for variant association testing. <sup>a</sup>Age reported as count, median (min-max).

### Hormone assays

T, DHEAS, sex hormone binding globulin (SHBG), luteinizing hormone (LH), follicle-stimulating hormone (FSH), and insulin levels were measured as previously reported^6^.

### Whole-genome sequencing (WGS)

Genomic DNA was isolated from whole blood samples using the Gentra Puregene Blood Kit (Qiagen, Valencia, CA). Whole-genome sequencing was performed by Complete Genomics, Inc., (CGI) using their proprietary sequencing technology. Their sequencing platform employed high-density nanoarrays populated with amplified DNA clusters called DNA nanoballs. DNA was read using a novel, iterative hybridization and ligation approach, which produces paired-end reads up to 70 bases in length^33^. CGI’s sequencing service included sample quality control, library construction, whole genome DNA sequencing, and variant calling. Reads were mapped to the NCBI Build 37.2 reference genome (GRCh37).

### Variant calling

Variant calling was performed using CGI’s Assembly Pipeline version 2.0. Although raw read data were provided, because of the unique gapped read structure produced by CGI sequencing, the use of other mapping or variant-call software was not recommended by CGI. Results were provided in CGI’s unique variant-call format. Variants included single nucleotide variants (SNVs), as well as insertions, deletions, and substitutions ≤50 bases in length. Calls were assigned confidence scores assuming equal allele fractions for the diploid genome (*varScoreEAF*). A Bayesian probability model was used to evaluate potential locus calls. The model accounted for read depth, base call quality scores, mapping/alignment probabilities, and empirical priors on gap sizes and discordance rates^34^. Based on the relative allele likelihoods a quality score was assigned for the chosen call:

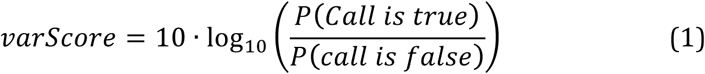

Scores were therefore reported in decibels (dB), where a score of 10dB represents a likelihood ratio of 10:1, 20dB means 100:1 likelihood, 30dB means 1000:1, etc. Quality thresholds for reporting variants were minimized (≥10dB for homozygous and ≥20dB for heterozygous variant calls) in order to maximize sensitivity. Variants were assigned a basic high/low quality flag (*VQHIGH* or *VQLOW*) based on a quality score threshold of 20dB for homozygous and 40dB for heterozygous variant calls.

### Called variant filtering

Variants were considered rare if they appeared with a frequency ≤2% in the Scripps Wellderly Genome Resource, which consisted of 597 unrelated participants of European Ancestry from the Scripps Wellderly Study^35^. The Wellderly study population is composed entirely of elderly individuals ≥80 years of age with no history of chronic disease. Within each sequenced family, reported variants that were inconsistent with Mendelian patterns inheritance were removed from consideration. A variant was considered consistent with Mendelian inheritance if it was called (*VQHIGH*) in one or more of the offspring and in at least one parent. The vast majority of DNA sequencing errors can be eliminated using Mendelian inheritance analysis^36^. As previously described^37–40^, an additional set of filters was applied to called variants for each sample in order to minimize the number of false positive calls: (i) variants with *VQLOW* allele tags were removed; (ii) variants in microsatellite regions were removed; (iii) variants within simple tandem repeat regions were removed; (iv) three or more SNPs clustered within a distance of 10bp were removed; (v) SNPs located within 10bp of an insertion or deletion (indel) were removed; (vi) calls located within known regions of segmental duplication were removed; (vii) calls with an observed read depth greater than 3× the average read depth (>168) were removed.

### Selection of optimal read depth and quality score thresholds

After systematically applying the filter matrix outlined above, optimal read depth and quality score thresholds were determined for each variant type by comparing calls between replicated samples for a particular family that was sequenced twice by CGI. Variants that were concordant between offspring sample pairs—above a given coverage depth and quality score threshold—were considered as true positive (TP) calls, while those that were discordant between replicates were considered false positives (FP). Accordingly, concordant variant calls that fell below a given depth and quality threshold were classified as false negative (FN) and discordant calls that fell below a given depth and quality threshold were classified as true negative (TN). Optimal depth and quality score thresholds were determined by selecting the thresholds that yielded the greatest Matthews correlation coefficient (MCC) values across each variant type^37^:

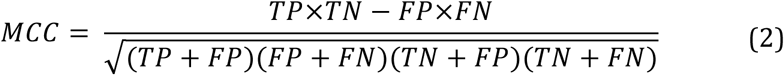

The thresholds chosen for association testing corresponded with the greatest MCC values for each variant type (**Table S2**).

### Assessing deleteriousness

After filtering for rare variants called with high confidence, as described above, variants were further characterized according to their predicted effects. Only variants that were predicted to have a deleterious effect were included in the association testing. Deleteriousness was primarily assessed using the Combined Annotation Dependent Depletion (CADD) tool^41^, which is trained on output from numerous annotation programs to predict deleteriousness based on conservation relative to our ancestral genome. Variants were retained if they produced a CADD score greater than the gene-specific mutation significance cutoff (MSC; 95% confidence interval) suggested by Itan *et al*.^42^. For variants outside of gene regions or in inconsistently annotated genes, the mean MSC (MSC_μ_ = 12.889) was used as the cutoff. In a further effort to reduce false positives, only coding variants classified as at least possibly damaging by the PolyPhen2^43^ or SIFT^44^ variant effect prediction tools and noncoding variants with LINSIGHT^45^ scores ≥0.8 were included in our analysis, as integrating individual methods can improve variant effect prediction^42, 46–48^. CADD and LINSIGHT are both primarily based on evolutionary conservation, and therefore carry certain limitations^49^. However, the applicability of prediction tools based on functional annotations from specific cell types^50, 51^ is extremely limited for PCOS because its pathophysiology involves numerous cell types across multiple organs^52^ and annotations for relevant cell types are largely unavailable^53–56^.

### Genome-wide rare variant association testing

Variants were then grouped for association testing using both gene-based and sliding window approaches. Gene regions included all coding and noncoding DNA from the 3’ UTR to 7.5kb upstream of the 5’ transcriptional start site (TSS) for all RefSeq annotated genes. The sliding window approach included all noncoding variation contained within sequential windows across the genome using three different window sizes: 10kb windows with no overlap, 25kb windows with 12.5kb overlap, and 100kb windows with 75kb overlap.

In rare variant association testing, genes are often filtered according to a minimum number of rare variants detected per gene^57–60^ or a minimum cumulative variant frequency (CVF) per gene^61–64^ in order to increase power to detect disease associations. In this study, genes and windows were removed from consideration if they did not harbor deleterious variants in at least 10% of affected subjects^65, 66^. For the gene-based test, in order to reduce the resultant bias towards larger genes, the observed CVFs were adjusted for gene length. Coding and noncoding CVFs were modeled separately using linear regressions against the coding sequence length^67, 68^ and the square root of noncoding sequence length^69, 70^, respectively. The models also accounted for gene-level GC content. The CVFs observed in affected subjects for each gene were then adjusted accordingly prior to applying the 10% threshold. Adjusting for gene length in this manner reduced the risk for erroneous associations, as genes with longer open reading frames are more likely to be reported falsely as disease-associated^71^.

Sequence kernel association test (SKAT) statistics were computed using the methods described in Schaid *et al.* ^72^, which account for pedigree information in calculating trait associations. Custom variant weights were applied for each variant according to their CADD Phred scores:

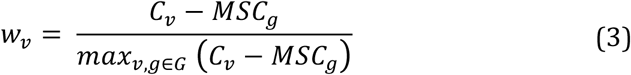

where *C_v_* is the CADD Phred score for variant *v*, *g* is the gene or window, *MSC_G_* is the mutation significance threshold for *g*, and *G* is the set of all variants within the genome. In this way, variants that are more likely to be deleteriousness were weighted more heavily than variants where the functional consequences are less likely to be damaging. For the dichotomous trait analysis, the binary outcome was adjusted for age and body mass index (BMI) using a logistic regression.

Six quantitative traits, T, DHEAS, LH, FSH, SHBG, and insulin levels, were tested for association. The trait distributions were each positively skewed. Testing against skewed trait distributions can result in heavily inflated Type I error rates in rare variant association testing^73^. Therefore, each trait was modeled against a gamma distribution when adjusting for age and BMI, and non-normal residuals (Shapiro-Wilk < 0.05) were then further normalized using a rank-based inverse normal transformation (INT). The INT has been found to be the optimal method for maintaining Type I error control without sacrificing power in rare variant association testing on non-normally distributed traits^74^.

### Quantitative trait meta-analysis

For complex diseases with multivariate phenotypes, combining multiple related phenotypic traits into one analysis can increase power in finding disease associations^75, 76^. We combined the six aforementioned quantitative trait associations into one meta-statistic using a Fisher combination function modified to account for correlated traits^77^. Inter-trait correlations were determined using the Pearson correlation coefficient (**Fig. S4**). P-values were adjusted using Bonferroni correction according to the number of variant groupings that were tested that contained at least one variant, as well as by the genomic inflation factor, λ. Correlation between meta-analysis association results and dichotomous trait association results were calculated using Spearman’s coefficient.

The characteristic disturbance of gonadotropin secretion associated with PCOS is increased LH relative to FSH release^78^. For genes with significant meta-analysis associations, LH:FSH ratios were compared between variant carriers and non-carriers using a Wilcoxon’s rank sum test, adjusted for multiple testing (Bonferroni). Differences in LH:FSH ratios between variant carriers and non-carriers would indicate that the gene variants alter gonadotropin signaling.

### *In silico* binding effect prediction

To assess the potential functional effects of noncoding variants identified in the quantitative trait meta-analysis, we predicted the corresponding impacts to transcription factor (TF) and RNA-binding protein (RBP) binding *in silico*. For each noncoding SNV identified in the quantitative trait meta-analysis, transcription factor (TF) binding affinities were calculated for all subsequences overlapping the SNV position on each strand within a ±20bp window. Binding affinity scores were calculated using position weight matrices (PWMs) derived from ENCODE ChIP-Seq experiments^71^. Scores were determined by summing the logged frequencies for a given sequence across a motif PWM. Binding p-values were defined as the probability that a sequence sampled from a genomic background distribution had an affinity score greater than or equal the largest affinity score produced from one of the tested subsequences. Genomic background sequences were generated using a first order Markov model^79^. The significance of a given change in binding affinity scores between reference and SNV alleles was assessed by determining whether the differences in relative binding affinity rank between the two alleles was significantly different than what would be expected by chance^80, 81^. P-values were conservatively adjusted to account for multiple testing using the Benjamini-Hochberg (BH) procedure^82^.

Once binding affinities were calculated for each TF at each SNV, filters were applied to identify the most likely candidates for TF binding site disruption. Instances in which the predicted TF binding affinity score was <80% of the maximum affinity score for the given motif were excluded. SNVs in which the reference allele was not predicted to bind a particular TF with statistical significance (P_BH_<0.05) were also removed from consideration, as well as variants in which both the reference and SNV alleles were predicted to bind a TF with statistical significance. Only TFs expressed in the ovary were analyzed. Tissue-specific gene expression was determined using GTEx data^83^ (median Reads Per Kilobase of transcript per Million mapped reads [RPKM] ≥0.1).

The identified SNVs were likewise analyzed for potential alteration to RNA-binding protein (RBP) sites following the same procedure but for a few modifications. RBPs and their binding affinity scores were determined using the ATtTRACT database^84^. Only sequences on the coding strand were evaluated as to reflect the mRNA sequences. Additionally, significant changes in RBP binding were considered regardless of the direction of effect, such that instances in which the alternate allele was predicted to induce RBP binding were also included.

### Supplementary Methods

For additional details regarding methodological considerations and rationale, please refer to the Appendix.

## RESULTS

### Characteristics of study population

The characteristics of the study population, including counts and trait distributions by familial relation and the numbers of subjects included in each association test, are summarized in **Table 1**.

### Whole-genome sequencing and variant calling

Genome sequencing yielded average genome coverage of 96.2% per sample and an overall mean sequencing depth of 56× (**Fig. S1**). On average, 90.4% and 66.4% of the genome, including 95.2% and 78.8% of the exome, was covered with at least 20× and 40× sequencing depth, respectively. Approximately 4.04 million high-confidence small variant calls were reported per genome. By applying optimal read depth and quality thresholds as well as a series of genomic filters^37^, we reduced the discrepancy rate between replicate samples from 0.23% to 0.04% for rare variants (**Tables S1 and S2**).

### Association testing and quantitative trait meta-analysis

We found 339 genes that had rare, deleterious variants in at least 10% of cases after adjusting for gene length and %GC content. No set of rare variants reached genome-wide significance for association with PCOS/HA disease status in the dichotomous trait analysis. We found 32 rare variants (2 coding, 30 noncoding) in the *DENND1A* gene that were collectively significantly associated with quantitative trait levels (*P*=5.31×10^−5^, *P_adj_*=0.019; **Table 2**), after adjusting for multiple testing and for observed genomic inflation (**Fig. S2**). Women with one or more of these *DENND1A* variants had significantly higher LH:FSH ratios (*P*=0.0012). PCOS/HA phenotype women with one or more *DENND1A* variants had significantly higher LH:FSH ratios than PCOS/HA phenotype women without *DENND1A* variants (*P*=0.0060). Unaffected women with one or more *DENND1A* variants had higher LH:FSH ratios than unaffected women without *DENND1A* variants (*P*=0.0586; **Fig. 1**), but the difference was not significant after multiple test correction (*P*<0.0167). No other gene-based set of rare variants reached genome-wide significance for association with quantitative trait levels. The correlation between the meta-analysis gene associations and the dichotomous trait gene associations was 0.24 (Spearman).

**Table 2.**
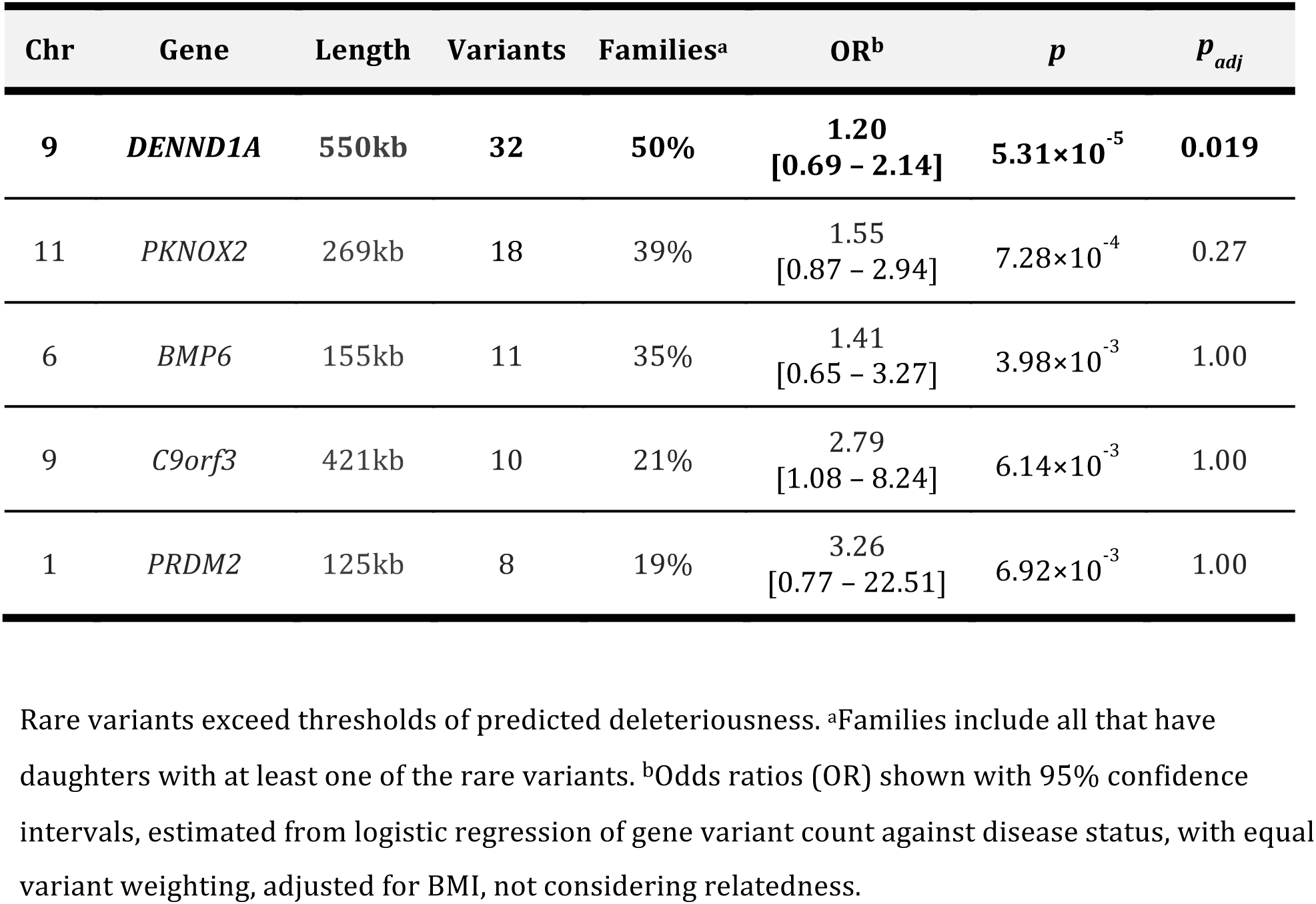
Top 5 rare variant associations from quantitative trait meta-analysis.

**Figure 1.**
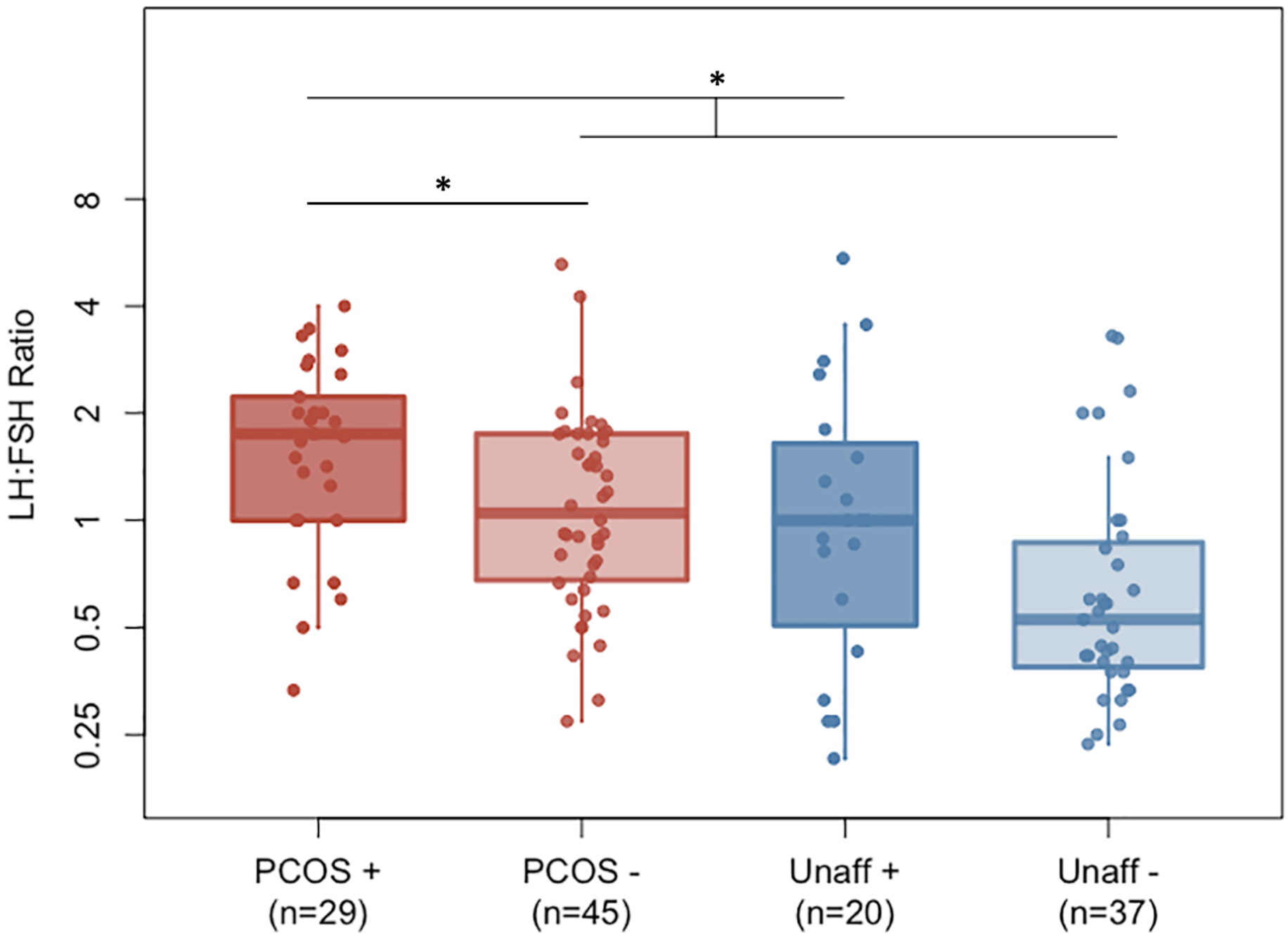
LH:FSH ratios in *DENND1A* variant carriers. LH:FSH ratios in DENND1A rare variant carriers (+) and non-carriers (−) in unaffected women and in women with PCOS/HA. Differences in group means were analyzed using Wilcoxon’s rank sum test (* P ≤ 0.0167).

Using the sliding windows approach, we found a subset of noncoding variants within a 25kb region of *DENND1A* (chr9:126,537,500-126,562,500) that were significantly associated with altered quantitative trait levels (*P*=1.92×10^−5^, *P_adj_*=9.53×10^−3^; **Fig. S3**; **Supplementary Data**). The region included three noncoding variants that were collectively present in eight PCOS/HA subjects and zero unaffected subjects. One of these variants, rs117893097 (MAF_Wellderly_ = 0.013), was homozygous in one of the subjects. This 25kb region encompasses one of the *DENND1A* GWAS risk variants (rs10986105; MAFWellderly = 0.034; OR_Meta_=1.39^85^), although none of the subjects with one of the rare variants in the region also had the rs10986105 risk allele. The relative positions of all of the rare variants found in *DENND1A* are shown in **Fig. 2**. No other windows across the genome were found to have significant associations with disease state or hormonal levels.

**Figure 2.**
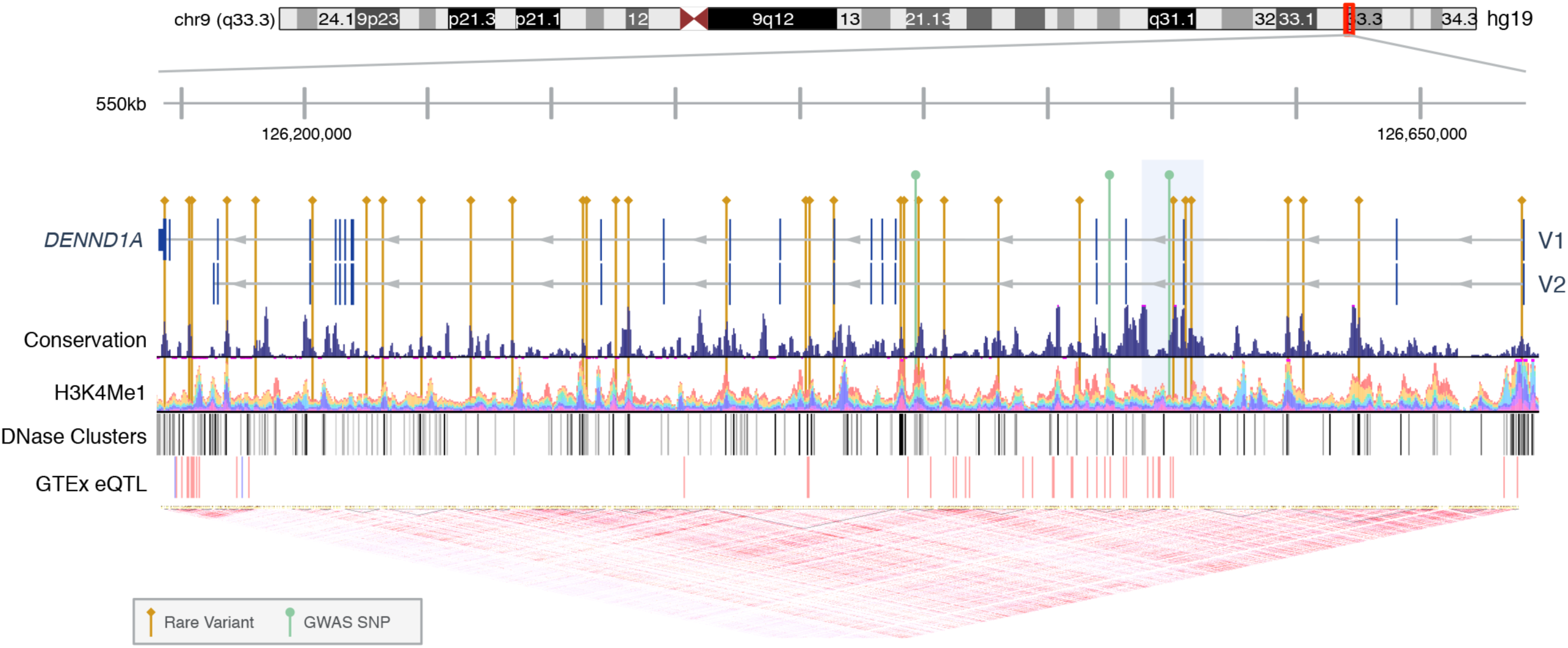
Rare Variants in *DENND1A*. The locations of deleterious rare variants and previously reported GWAS SNPs within the DENND1A gene, including the two primary isoforms DENND1A.V1 and DENND1A.V2. The 25kb region significantly associated with altered hormone levels is highlighted in light blue. The Conservation track was measured on multiple alignments of 100 vertebrate species by phyloP^113^. The H3K4Me1 track shows enrichment of mono-methylation of lysine 4 of the H3 histone protein, which is associated with enhancers and DNA regions downstream of transcription starts, as determined by ChIP-seq assay and layered by different cell types^53^. The DNase Clusters track shows regions of DNase hypersensitivity, an indicator of regulatory activity, with darkness proportional to maximum signal strength53. The GTEx eQTL track displays gene expression quantitative trait loci for DENND1A, as identified from GTEx RNA-seq and genotype data, with red and blue indicating positive and negative effects on gene expression, respectively^114^. The linkage disequilibrium heatmap was generated using Phase1 CEU data from the 1000 Genomes Project^115^.

Several other PCOS GWAS candidate genes appeared in our filtered set of genes, including *C9orf3*^5, 6, 8^ (*P*=6.14×10^−3^), *HMGA2*^5^ (*P*=0.062), *ZBTB16*^8^ (*P*=0.20), *TOX3*^5, 8^ (*P*=0.22), and *THADA*^4, 5, 7, 8^ (*P*=0.74). *C9orf3* had the 4^th^ strongest association overall (**Supplementary Data**), but failed to reach genome-wide significance after correction for multiple testing. The relative quantitative trait associations for these genes are illustrated in **Fig. 3**.

**Figure 3.**
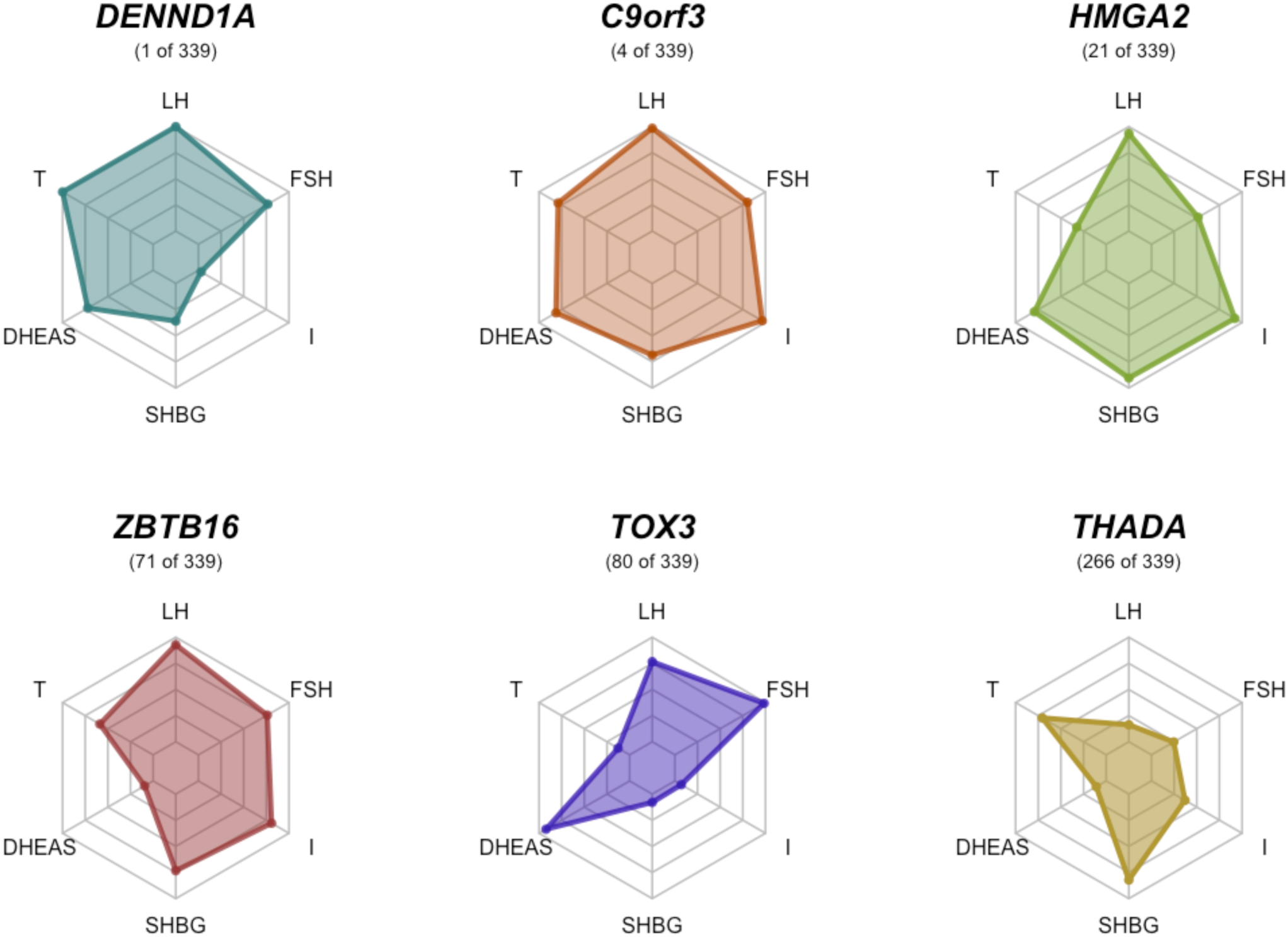
Trait associations for PCOS GWAS genes. The relative quantitative trait associations are shown for PCOS GWAS susceptibility loci included in meta-analysis results, with meta-analysis association ranking.

### Protein binding effect prediction

Nine of the *DENND1A* variants were predicted to significantly impact TF binding motifs, while the majority of variants were predicted to significantly alter RBP binding motifs (17 disrupted, 14 induced). The specific TF and RBP motifs associated with each noncoding variant are listed in **Table 3**. Binding by the heterogeneous nuclear ribonucleoprotein (hnRNP) family of RBPs appeared to be the most commonly impacted.

**Table 3.**
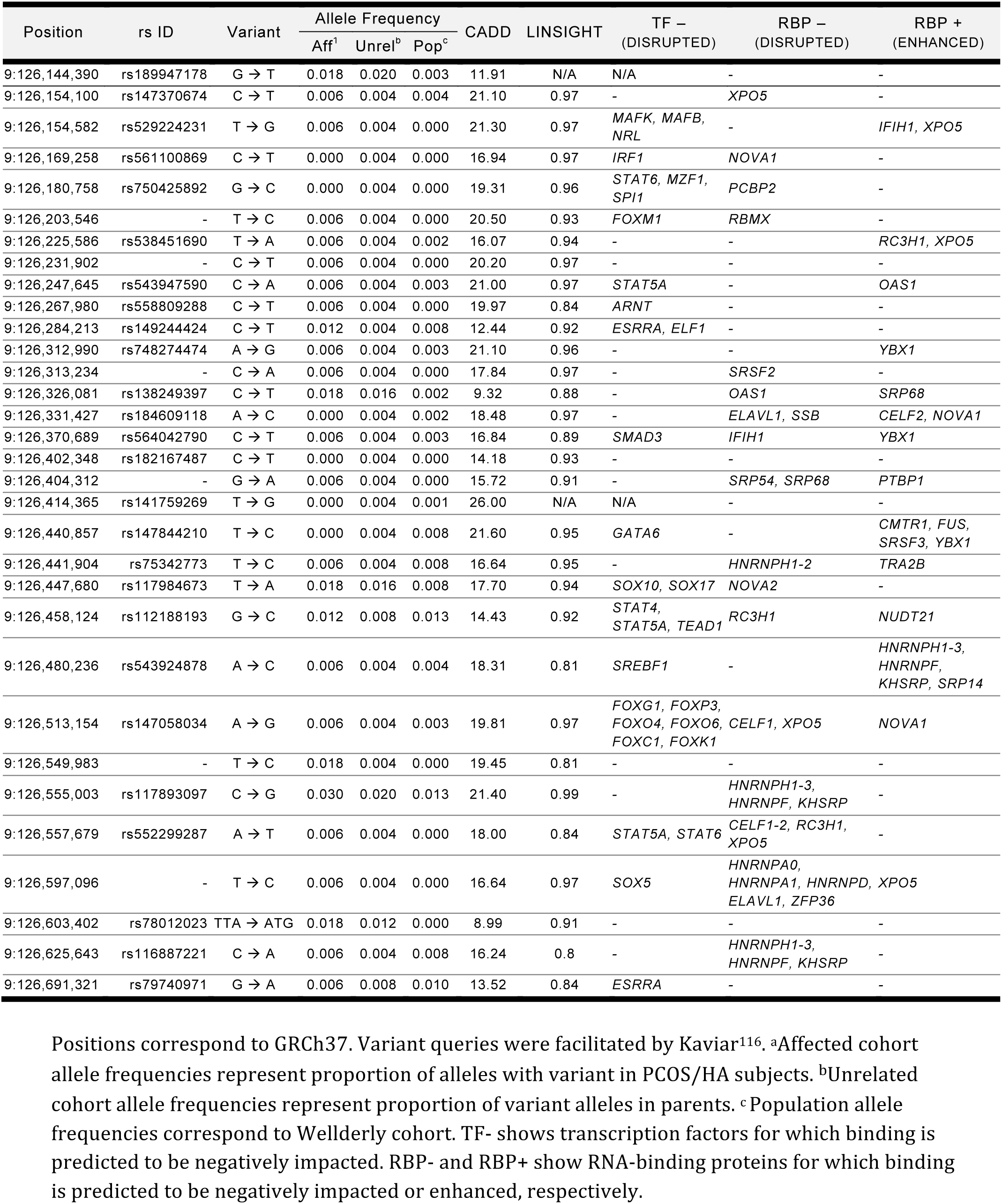
Deleterious, rare variants in DENND1A.

## DISCUSSION

We identified rare variants in *DENND1A* that were significantly associated with altered reproductive and metabolic hormone levels in PCOS. These findings are of considerable interest because common SNVs in *DENND1A* were associated with PCOS diagnosis in a GWAS of Han Chinese women^4^; these associations were subsequently replicated in women of European ancestry^86^. Our study, using an independent family-based WGS analytical approach, provides further evidence that *DENND1A* is an important gene in the pathogenesis of PCOS. These findings complement the studies of McAllister and colleagues^56, 87^ that have shown that *DENND1A* plays a key role in androgen biosynthesis in human theca cells and is upregulated in PCOS theca cells.

*DENND1A* encodes a protein that is a member of the connecdenn family of proteins, which function as guanine nucleotide exchange factors for the Rab family of small GTPases^77^. The DENND1A protein, also known as Connecdenn 1, is thought to link Rab35 activation with clathrin-mediated endocytosis^42^. Following its reported associations with PCOS^4, 86^, McAllister and colleagues^87^ investigated the role of *DENND1A* in ovarian androgen biosynthesis, a key biologic pathway that is disrupted in PCOS^88^. *DENND1A* encodes two transcripts as the result of alternative splicing, DENND1A.V1 and DENND1A.V2^56^. The encoded V2 protein was found in ovarian theca cells and its abundance was correlated with increased androgen production^87^. The expression of V2 was increased in PCOS theca cells^87^. Forced expression of the *V2* transcript produced a PCOS phenotype in normal theca cells, whereas knockdown of *V2* in PCOS theca cells reduced thecal androgen biosynthesis^87^. Urine exosomal V2 mRNA was also increased in PCOS women^87^. Taken together, these findings provide strong support for the hypothesis that *DENND1A* plays a role in PCOS pathogenesis. The increased LH:FSH ratios that we observed in the *DENND1A* rare variant carriers (**Fig. 1**) suggest that *DENND1A* plays a role in the regulation of gonadotropin secretion^89^.

The *DENND1A* risk variants identified by GWAS are located in introns and the functional consequences of these variants are unknown^56^. There have been no large-scale sequencing studies reported to map causal variants in *DENND1A* ^85^. Targeted^87, 90^ and whole exome sequencing^91^ in small cohorts of PCOS women have failed to identify any coding variants in *DENND1A* that were associated with PCOS or with V2 isoform expression. Genomic sequencing of the intronic region where V1 and V2 are alternatively spliced also failed to identify any variants that consistently favored V2 expression in a study of 20 normal and 19 PCOS women^56^. The GWAS risk variants and most of the variants identified in our study lie well upstream (100-400kb) of this region (**Fig. 2**).

Many of the *DENND1A* variants identified in the present study were predicted to disrupt conserved TF binding motifs, which could affect gene expression, but most of the variants were predicted to alter affinities of RBPs to the mRNA transcript (**Table 3**). It is plausible, therefore, that the rare variants we reported were selectively driving the expression of the V2 splicing variant via post-transcriptional regulation. Collectively, the *DENND1A* variants identified in this study were found in 50% of families, but each individual variant was typically found in only one or two families. Our findings, therefore, support a model of PCOS in which causal variants are individually uncommon but collectively tend to occur in key genes. Our recent findings^92^ of multiple rare exonic rare variants in the *AMH* gene that reduce its biologic activity in ∼3% of women with PCOS is consistent with this model.

Our results also align with the emerging evidence that rare coding variants with large effect sizes do not play a major role in complex disease^93^. Rather, it appears that complex traits are primarily driven by noncoding variation^94, 95^, both common and rare^96,97^. Of the 32 rare variants predicted as deleterious that we identified in *DENND1A*, 30 were noncoding. Rare variant association studies typically require very large sample sizes^98^, but paired WGS and transcriptome sequencing analysis from one large family demonstrated that rare noncoding variants have strong effects on individual gene-expression profiles^99^. In a similarly designed study to ours, Ament and colleagues^100^ identified rare variants associated with increased risk of bipolar disorder, the vast majority of which were noncoding. Due to the relative cost-effectiveness of whole exome sequencing^94^, the limited availability of computational tools designed to predict the effects of noncoding variants on phenotypes^101^, and our relatively poor understanding of regulatory mechanisms in the genome^102^, noncoding variants have been noticeably understudied in complex trait genetics. As larger WGS datasets are accumulated and the focus of complex trait studies shifts more towards understanding regulatory mechanisms, the contribution of rare noncoding variants in various complex diseases will become clearer.

Several established PCOS candidate genes besides *DENND1A* appeared among the top gene associations, but failed to reach genome-wide significance. These genes included previously reported PCOS GWAS susceptibility loci, *C9orf3*, *HMGA2*, *ZBTB16*, *TOX3, and THADA* (**Fig. 3**). Two additional genes with strong, but not genome-wide-significant, associations with PCOS quantitative traits are highly plausible PCOS candidate genes. *BMP6* had the third strongest association in our meta-analysis (*P*=4.00×10^−3^, **Table 2**). It is a member of the bone morphogenetic protein family, which are growth factors involved in folliculogenesis. *BMP6* expression was previously found to be significantly higher in granulosa cells from PCOS women compared with reproductively normal control women^103^. Moreover, BMP6 was found to increase expression of the FSH receptor, inhibin/activin β subunits, and AMH genes in human granulosa cells^104^. *PRDM2* had the fifth strongest association in our meta-analysis (*P*=6.95×10^−3^, **Table 2**). *PRDM2* is an estrogen receptor co-activator^105^ that is highly expressed in the ovary and pituitary gland^83^. Ligand bound estrogen receptor alpha (ERα) binds with PRDM2 to open chromatin at ERα target genes^105,106^. PRDM2 also binds with the retinoblastoma protein^107^, which has been shown to play an important role in follicular development in granulosa cells^108,109^.

The central statistical challenge in studying rare variants is achieving adequate power to detect significant associations while controlling for Type I error. The family-based structure of our cohort provided an enrichment of individual rare variants and enabled modeling of familial segregation^110^. To mitigate variant calling errors, we utilized replicate samples from one family to determine optimal read depth and quality thresholds. To remove irrelevant variants from consideration, we applied a LINSIGHT score threshold and gene-specific CADD score thresholds^72^ to filter for deleteriousness. We further prioritized variants by weighting them by their relative CADD scores. To group rare variants effectively, we applied several windows-based binning methods, in addition to the gene-based approach, to ensure that different kinds of functionally-correlated genomic regions were tested, both of fixed length and of variable length. To limit our search to genes that were more likely to have specific roles in PCOS etiology, we only considered genes with rare deleterious variants in at least 10% of cases, as causal rare variants are more likely to accumulate in core disease genes^94^. We greatly increased our power to detect relevant disease genes and account for pleiotropic effects by consolidating quantitative trait association results into a meta-analysis^75, 76^. In order to reduce Type I error, in addition to the variant calling quality control measures, we modeled the quantitative traits against skewed distributions and further normalized trait residuals using an INT.

Given the size of our cohort, it was necessary to apply relatively strict filters based on predicted variant effects and cumulative allele frequencies in order to detect rare variant associations. Very large sample sizes are otherwise required for WGS studies of rare variants^98^. By only including genes with rare, likely-deleterious variants in at least 10% of cases, we greatly reduced the multiple-testing burden of our analysis, thereby increasing our power to detect core PCOS genes^111, 112^. Applying *a priori* hypotheses regarding which variants and genes may be relevant to disease, however, is analogous to a candidate gene approach^98^. Any sets of rare variants that contribute to PCOS in smaller subpopulations of PCOS women, as we found for *AMH*^92^, were likely removed from consideration. Likewise, any causal rare variants that were not predicted bioinformatically to be deleterious based on existing annotations and evolutionary conservation^41, 45^ would not have been detected. Furthermore, because variants were filtered for consistency with Mendelian inheritance, *de novo* mutations were not considered.

Because the number of genetic variants is directly correlated with the size of the gene^71^, the CVF threshold introduced a bias towards larger genes in some of the analyses. However, this bias was mitigated by adjusting for gene length. Furthermore, any such bias was not applicable to our fixed-length windows-based approach, which replicated our *DENND1A* findings. It is also possible that causal rare variants with larger effect sizes were omitted from the meta-analysis because we tested against normalized trait residuals in an effort to reduce Type I errors. Using normalized trait residuals may have excluded variants with large effects that produced outliers. However, as mentioned above, recent evidence has demonstrated that complex traits are primarily driven by noncoding variation with modest effect sizes^94, 95^.

Despite the numerous steps taken to increase power, our study ultimately remained underpowered to detect rare variant associations with PCOS diagnosis in the dichotomous trait analysis. Although an association with PCOS quantitative traits implicates genetic variants in disease pathogenesis, it does not necessarily mean that a gene is associated with PCOS itself. The correlation between the dichotomous trait results and quantitative trait meta-analysis results was 0.24. The noncoding variants identified in this study, despite being rare and predicted to be deleterious, may be in linkage disequilibrium with the actual pathogenic variant on the same alleles that were removed by the applied allele frequency or predicted effect thresholds. Replication and functional studies are needed to confirm individual variant functionality and disease associations.

In summary, by applying family-based sequence kernel association tests on filtered whole-genome variant call data from a cohort of PCOS families, we were able to identify rare variants in the *DENND1A* gene that were associated with quantitative hormonal traits of PCOS. Our results suggest that rare noncoding variants contribute to the distinctive hormonal profile of PCOS. This study also demonstrates that using a quantitative trait meta-analysis can be a powerful approach in rare variant association testing, particularly for complex diseases with pleiotropic etiologies.

## ACKNOWLEDGEMENTS

This study was supported by US National Institutes of Health (NIH) grants P50 HD044405 (A.D.) and R01 HD085227 (A.D.). M.D. was supported by NRSA fellowship T32 DK007169. This study uses data from the Scripps Wellderly Genome Resource, which is funded under NIH grant 5 UL1 TR001114 Scripps Translational Science Institute CTSA Award. We thank Drs. Terry Farrah and Gustavo Glusman, from the Institute for Systems Biology for facilitating variant queries via Kaviar (http://db.systemsbiology.net/kaviar). We thank the Northwestern University Research Computing Services team for supporting the computational needs of this research. We are very grateful to all of the families who took part in the study.

## AUTHOR CONTRIBUTIONS

M.D. M.U., A.D, and M.G.H. conceived and designed the study. M.D. performed experiments and statistical analyses. R.S. aided with data management, project coordination, and statistical analyses. R.S.L. and A.D. recruited study subjects and measured or analyzed phenotypic data. M.D. wrote the paper. All authors critically reviewed and approved the paper.

